# Indirect assortative mating for human disease and longevity

**DOI:** 10.1101/185207

**Authors:** Konrad Rawlik, Oriol Canela-Xandri, Albert Tenesa

## Abstract

Phenotypic correlations of couples for phenotypes evident at the time of mate choice, like height, are well documented. Similarly, phenotypic correlations among partners for traits not directly observable at the time of mate choice, like longevity or late-onset disease status, have been reported. Partner correlations for longevity and late-onset disease are comparable in magnitude to correlations in 1^st^ degree relatives. These correlations could arise as a consequence of convergence after mate choice, due to initial assortment on observable correlates of one or more risk factors (e.g. BMI), referred to as indirect assortative mating, or both. Using couples from the UK Biobank cohort, we show that longevity and disease history of the parents of white British couples is correlated. The correlations in parental longevity are replicated in the FamiLinx cohort. These correlations exceed what would be expected due to variations in lifespan based on year and location of birth. This suggests the presence of assortment on factors correlated with disease and lifespan, which show correlations across generations. Birth year, birth location, Townsend Deprivation Index, height, waist to hip ratio, BMI and smoking history of UK Biobank couples explained ~70% of the couple correlation in parental lifespan. For cardiovascular diseases, in particular hypertension, we find significant correlations in genetic values among partners, which support a model where partners assort for risk factors genetically correlated with cardiovascular disease. Identifying the factors that mediate indirect assortment on longevity and human disease risk will help to unravel what factors affect human disease and ultimately longevity.

## Background

Partner correlations for a variety of phenotypes have been reported when examining environmental and genetic contributions to complex traits (1-11). These correlations between nominally unrelated individuals are substantial, with magnitude comparable to correlations between first degree blood relatives, for instance, between parents and children (9, 10). Such effects can be interpreted as phenotypic convergence among partners due to the environmental factors that partners share during their co-habitation. In the case of late-onset diseases and longevity, which are not directly observable or present at the time of mate choice, this would arguably be the simpler explanation. Alternatively, partner correlations for late onset disease and longevity could arise due to indirect assortative mating. That is, direct assortative mating for traits, characteristics or social factors that are risk factors of disease and potentially observable at the time partners met (for instance, behavioural risk factors of disease such as smoking) would lead to indirect assortative mating for other focal traits, such as longevity or late-onset disease. The distinction between the causes that underpin partner effects has implications for the study of human behaviour, epidemiology and population genetics. It provides information about human mate choice behaviour and informs about the importance of environmental risk factors shared by couples in the household. The importance to population genetics arises because assortative mating for heritable traits induces a correlation of genetic values among partners, whilst assortment on environmental factors (e.g., social homogamy), and environmental effects shared by partner do not. The correlation of the genetic values of the partners in turn affect the amount of genetic variance of the trait assorted on, as a consequence estimates of heritability reported in the literature which do not account for assortment overestimate the heritability for that trait in a random mating population due to the covariance among alleles at different loci (12) (Fig. 1a, Methods). Furthermore, assortative mating for a trait would also induce an increase in heritability for genetically correlated traits (13) (Fig. 1b) and a change in the genetic correlation between the assortment and focal traits (Fig. 1c). This is the case even if these focal traits do not directly underlie mate choice, or do not manifest at the time of mate choice. For instance, assortment for BMI, would induce an indirect increase in the genetic variance of cardiovascular disease because there is a positive genetic correlation between these two traits (14), and an increase in their genetic correlation with respect to what would be expected under random mating.

**Figure 1:**
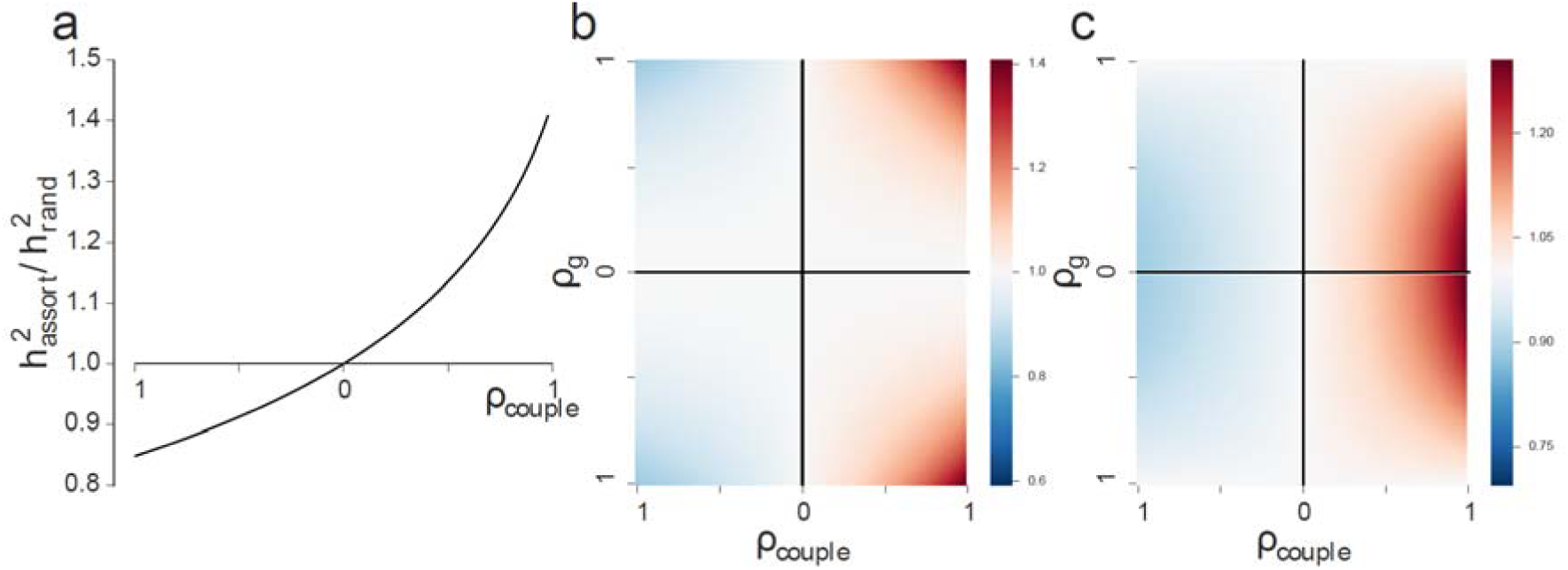
Effects of indirect assortative mating on heritability and correlations. We consider a pair of traits (Methods), a trait which is the target of assortment, e.g., BMI, and a genetically correlated focal trait, e.g., hypertension disease liability, both with heritabilities of 0.3 in a random mating population. We illustrate relative changes in heritability of the assortment trait (**a**), heritability of the focal trait (**b**) and genetic correlation between the traits (**c**) as functions of the strength of assortative mating (ρ_couple_) and genetic correlation in a random mating population between the traits (ρ_g_). Specifically in all three panels we plot the ratios of the parameter under assortment to random mating. We assume a population at equilibrium after assortative mating (which happens only after a few generations of assortment (13)) relative to a random mating population. In (**b**) and (**c**) red colors indicate areas where assortative mating leads to increased genetic variance in the focal trait and increased absolute genetic correlations, i.e., the ratio of 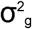 or ρ_g_ after assortative mating to that in a random mating population is greater than one.

Here, we present data showing that there is indirect assortment for both longevity and risk of disease. Specifically we find that humans choose partners with similar parental history of disease and parental longevity. Since partner choice most likely happens before the parental onset of most of these diseases or parental death, these are unlikely to be the traits on which such choice is made. Furthermore as these traits are heritable indirect assortment present the most parsimonious model. Finally, we demonstrate assortment directly, showing that the genetic values (i.e. polygenic risk scores) for hypertension are correlated among partners. Given that assortment for hypertension itself is unlikely, we hypothesise that this correlation in genetic values arises through assortment for one or more traits that influence mate choice and which are genetically correlated with hypertension.

## Results and discussion

Partner correlations for age at death have been demonstrated going back to early work on assortative mating (1). Similarly, we found that the ages of death of the biological mothers and fathers of all self-reported White-British individuals in the UK Biobank with both parents deceased (N=252,899 pairs of parents of UK Biobank participants) was significantly different from zero (ρ =0.11, pval<10^−188^). This correlation was only slightly reduced (ρ=0.10) and still highly significant (pval<10^−188^) when adjusting for the participants’ year of birth as a proxy of the parent’s year of birth, which itself was unavailable. We replicated this finding the FamiLinx (15) cohort. Partner correlations for longevity in 239,541 couples of individuals born across the world between 1600 and 1910 in the FamiLinx (15) cohort were significantly higher (ρ=0.18, pval<10^−188^) which is expected due to the broader range of birth years and wider geographical distribution of this cohort. After adjusting for sex, birth year and birth place (Methods), partner correlations for longevity were ρ=0.13 (pval<10^−188^), which, although slightly higher, is comparable to those in the UK Biobank cohort. These correlations are significantly lower than the correlation of 0.23 reported a century ago for a much smaller sample from the UK (1), but similar to more recent estimates of 0.12 in a Canadian population (16). Estimates of heritability for longevity in the FamiLinx cohort (15) imply a correlation between 1^st^ degree relatives of 0.06, while previous estimates of heritability suggest higher correlations of 0.13(17), suggesting that partner effects are comparable in magnitude, or even exceed, genetic effects on longevity. The age of death of partners could potentially be correlated due to effects directly related to the partner’s death (i.e. a partner’s death has a causal link with the other partner’s death), which together with the assortment by birth year would lead to partner correlations for lifespan. More generally, convergence due to shared environmental factors, represents in the absence of other data the most plausible explanation for the observed partner correlations. We therefore studied the correlation between partners in the lifespan of their parents for which no obvious direct link should exist. As longevity has a heritable component, the existence of such correlations in parental phenotypes would suggest the possibility that the observed partner correlations partially arise due to indirect assortment on heritable risk factors. Considering, from amongst 79,094 White-British couples among UK Biobank participants, the 40,504 and 60,978 couples with, respectively, both mothers and both fathers deceased, we found significant correlations for the lifespans of both the mothers (ρ=0.043, pval=10^−9^) and the fathers (ρ=0.025, pval=10^−5^) (Methods). Considering parents of couples in the FamiLinx(15) cohort, we again observed higher correlations in lifespans of mothers (ρ=0.061, pval=10^−55^) and fathers (ρ=0.071, pval=10^−107^) compared to the UK Biobank, although correlations between adjusted lifespans where again comparable to those in the UK Biobank (ρ=0.03, pval=10^−17^ and ρ=0.02, pval=10^−7^ for fathers and mothers respectively). The observed partner correlations in parent’s lifespans are expected to be partly explained by differences in life expectancy across history and geography. In order to confirm that they are not purely a consequence of assortment for year and place of birth we simulated alternative populations of couples maintaining the assortative mating structure for these factors (Methods). The observed correlations lie in the extreme tails of the respective distribution of correlations between parents’ lifespans in this fictitious mating structure (SI Appendix Figure S1), with empirical pvalues of 0.0002 and <0.0001 for mothers of couples in UK Biobank and FamiLinx respectively and 0.0093 and <0.0001 for the fathers of couples in UK Biobank and FamiLinx respectively, confirming that they are unlikely to be an artefact of the age or birth structure of the data. Year and birth place, socioeconomic status (as measured by Townsend Deprivation Index), height, waist to hip ration, BMI and smoking history measured in Pack Years (as a proxies of a putative behavioural factor associated with disease and longevity), show significant partner correlations (SI Appendix Table S1) in the UK Biobank and are some among all possible factors explaining longevity. We therefore examined the combined effect of these factors, on the observed correlations of longevity among the mothers and fathers of couples by evaluating the correlations in residuals of regressing parental longevity on these factors and, in the case of continuous factors, their squares (Methods and SI Appendix Table S2). Assortment for birth year and location were the most important factors, reducing the observed correlations for both maternal and paternal longevity by around 55%. Socioeconomic status and the other factors had a lesser but still important effect on the correlation of lifespan of parents, reducing such correlation an additional ~15%. This suggests these factors and socioeconomic status are correlated across generations as the children’s phenotypes and socioeconomic status explain some of the correlation in longevity of their respective parents. Using subsets of 79,216 and 64,002 genotyped unrelated White-British individuals in the UK Biobank with respectively deceased fathers and mothers, we estimated heritabilities and genetic variant effects for parental longevity based on common variants (MAF > 5%) (Methods). Significant heritabilities were observed for mothers (h^2^=0.03) and fathers (h^2^=0.04) (SI Appendix Table S3). We then estimated genetic values (18, 19) (i.e. Best Linear Predictors, BLUPs) for parental longevity, and used a subset of 10,160 genotyped White-British couples to estimate partner correlations in genetic effects. These were found not to be significantly different from zero (ρ= −0.007, pval = 0.6 and ρ=0.01, pval=0.3 for paternal and maternal longevity respectively). Polygenic risk scores for variants known to be associated with longevity were not significantly correlated among partners (ρ= 0.001, pval = 0.9 and ρ=-0.01, pval=0.1 for paternal and maternal longevity respectively). The lack of correlations in genetic values is consistent with environmental assortment. However, power to detect correlations in genetic values is limited due to the low number of couples available and the low heritability of the trait (SI Appendix Table S4).

We hypothesised that if the lifespan of the mothers and fathers of couples was correlated, then their disease history could also be correlated. Disease history for both biological parents of each partner was reported by 58,043 couples for Heart Disease, Stroke, Chronic Bronchitis, High Blood Pressure, Diabetes and Alzheimer’s Disease and by 57,644 couples in the case of Lung Cancer, Bowel Cancer, Parkinson’s Disease and Depression. For the latter subset, information regarding disease history for the relevant parent for Breast and Prostate Cancer was available for each partner. We found significant (P<0.05) polychoric correlations consistent for both fathers and mothers for half of the twelve diseases: heart disease, stroke, lung cancer, chronic bronchitis, hypertension, and Alzheimer’s disease (Table 1, SI Appendix Table S4), with only stroke in fathers failing significance after Bonferroni correction. Of these, the largest correlation was for paternal hypertension (ρ=0.09) and the smallest for paternal stroke (ρ=0.02). The history of prostate cancer among fathers of couples was also significantly correlated (ρ=0.07, pval=0.004). Among mothers, the correlations for lung cancer, hypertension and Alzheimer’s were the largest (ρ=0.08), whilst the correlations for chronic bronchitis and heart disease were only marginally smaller (ρ=0.06). In order to exclude the possibility of assortment on the individuals own disease status, we repeated the analysis using only couples were neither of the partners had reported the disease, i.e., were both self-reported as controls. This was largely in agreement with the analysis using all couples (SI Appendix Table S5). Furthermore, we confirmed the observed correlations are not purely a consequence of the mating structure due to year and location of birth employing the same permutation approach used for longevity (SI Appendix Table S6). Results for permutations using the parent’s year of birth, available in only a subset of parents, did not suggest that using the offspring’s year of birth as a proxy introduced a substantial bias (Methods). Using information from 114,264 unrelated genotyped White-British individuals, we estimated heritabilities (SI Appendix Table S7) and variant effects of the studied parental disease histories using common variants (MAF > 5%). We then computed genetic values for 10,160 genotyped White-British couples for both maternal and paternal family history of each of the diseases. Correlations among couples in genetic values would indicate that phenotypic partner correlations arise not only through common environment but also through assortative mating. Correlations between genetic values of partners were significant (pval < 0.05) for maternal and paternal history of hypertension as well as maternal heart disease, stroke and chronic bronchitis (Table 2) with only maternal chronic bronchitis and hypertension significant after Bonferroni correction. Whilst hypertension in fathers did not reached the stringent Bonferroni correction threshold, the size of the correlation was similar to that of maternal hypertension. Furthermore, hypertension remained significant in the meta-analysis of paternal and maternal correlations (Table 2). To assess whether the observed correlations could arise due to temporal or geographical stratification in the population, we recomputed SNP effects adjusting for Birth Year, Birth Location and the relevant parental age (i.e. either reported age or age at death). While correlations between genetic values were reduced, they remained significant (pval < 0.05) for maternal and paternal hypertension and maternal chronic bronchitis and stroke (SI Appendix Table S8). Finally, we repeated the previous analysis but now using own disease status instead of parental disease status. We restricted the analysis to diseases with prevalence in the sample above 5% and excluding prostate and breast cancers (Table 2). Despite the small sample size, we again find the correlations of genetic value of partners for hypertension to be significant and of similar size to the parental hypertension (ρ=0.03, pval = 0.005), thus indicating indirect genetic based assortment also for the UK Biobank participant’s own disease status. This correlation is likely indirectly generated through genetic correlation between the focal trait (i.e. hypertension) and an other, genetically correlated, trait or traits for which assortment happens, e.g., BMI (20).

**Table 1:** Polychroic correlations for family history

|  | Father |  |  | Mother |  |  |
| --- | --- | --- | --- | --- | --- | --- |
| | $\rho_{chor}$ | s.e. | $P$ | $\rho_{chor}$ | s.e. | $P$ |
| Heart Disease | 0.04 | 0.006 | $6 \times 10^{-11}$ | 0.07 | 0.007 | $9 \times 10^{-23}$ |
| Stroke | 0.02 | 0.009 | 0.003 | 0.06 | 0.009 | $2 \times 10^{-11}$ |
| Lung Cancer | 0.04 | 0.012 | $1 \times 10^{-4}$ | 0.08 | 0.018 | $1 \times 10^{-5}$ |
| Bowel Cancer | 0.04 | 0.015 | 0.009 | -0.01 | 0.017 | 0.747 |
| Breast Cancer | - | - | - | 0.01 | 0.012 | 0.325 |
| Chronic Bronchitis | 0.06 | 0.01 | $2 \times 10^{-9}$ | 0.06 | 0.015 | $7 \times 10^{-5}$ |
| High Blood Pressure | 0.09 | 0.007 | $1 \times 10^{-35}$ | 0.08 | 0.006 | $7 \times 10^{-38}$ |
| Diabetes | 0.02 | 0.012 | 0.067 | 0.04 | 0.011 | 0.001 |
| Alzheimer's | 0.07 | 0.017 | $2 \times 10^{-5}$ | 0.08 | 0.011 | $3 \times 10^{-13}$ |
| Parkinson's | 0.02 | 0.027 | 0.267 | 0.04 | 0.034 | 0.13 |
| Depression | 0.03 | 0.022 | 0.103 | 0.04 | 0.014 | 0.005 |
| Prostate Cancer | 0.04 | 0.013 | 0.004 | - | - | - |

$\rho_{chor}$ = polychoric correlation, s.e. = standard error, $P$ = pvalue for $\rho_{chor} = 0$

**Table 2:** Within couple correlations of genetic values for family history and self-reported disease.

|  | Father |  | Mother |  | Combined<sup>†</sup> |  | Self<sup>*</sup> |  |
| --- | --- | --- | --- | --- | --- | --- | --- | --- |
| | $\rho$ | $P$ | $\rho$ | $P$ | $\rho$ | $P$ | $\rho$ | $P$ |
| Heart disease | 0.013 | 0.18 | 0.02 | 0.05 | 0.016 | $9 \times 10^{-3}$ | -0.015 | 0.14 |
| Stroke | 0.002 | 0.85 | 0.024 | 0.01 | 0.013 | 0.12 | 0.004 | 0.7 |
| Lung cancer | -0.006 | 0.56 | 0.016 | 0.12 | 0.005 | 0.32 | - | - |
| Bowel cancer | -0.001 | 0.95 | -0.016 | 0.1 | -0.008 | 0.14 | - | - |
| Breast cancer | - | - | -0.004 | 0.68 | - | - | - | - |
| Chronic bronchitis | 0.006 | 0.52 | 0.032 | 0.001 | 0.019 | 0.07 | 0.011 | 0.26 |
| High blood pressure | 0.03 | 0.002 | 0.03 | 0.002 | 0.030 | $8 \times 10^{-6}$ | 0.028 | 0.005 |
| Diabetes | 0.009 | 0.37 | 0.01 | 0.32 | 0.009 | 0.09 | 0.024 | 0.02 |
| Alzheimer's | 0.001 | 0.9 | 0.007 | 0.45 | 0.004 | 0.27 | - | - |
| Parkinson's | -0.002 | 0.82 | -0.001 | 0.95 | -0.001 | 0.42 | - | - |
| Severe depression | 0.017 | 0.1 | -0.01 | 0.3 | 0.003 | 0.41 | 0.017 | 0.09 |
| Prostate cancer | 0.009 | 0.34 | - | - | - | - | - | - |

<sup>†</sup>meta-analysis of paternal and maternal results, <sup>\*</sup>contains only results for self-reported non sex specific disease
with UK Biobank prevalence > 5%, $\rho$ = Pearson's correlation between genetic values in couples, $P$ = pvalue for
$\rho=0$

## Conclusions

Taken together the results suggest that the characteristics that influence mate choice lead to detectable assortment for familial disease and longevity. This assortment is only partially explained by birth cohort and the few factors chosen to reflect the social mating structure, suggesting a contribution to assortment for parental disease history and longevity of other traits, lifestyle choices or social factors shared among parents and children. While we have not directly demonstrated that the underlying factors are transferred across generations, that is, that the same behavioural or social factors which drive parental disease risk are also the factors underlying mate choice in the offspring, such a model presents the most canonical explanation. Furthermore, the presence of correlations in genetic values for parental and maternal family history as well as self-reported status for hypertension provides direct evidence for the presence of assortment on heritable and genetically correlated risk factors for this disease. Two consequences of this model are that partner effects for longevity and disease are partly explained by indirect assortative mating and partly by shared environment, and that disease prevalence in the population may potentially be increased through indirect assortment for traits or risk factors correlated with disease (21).

## Methods

### Effect of assortative mating on genetic variance and genetic correlation

We compute effects of indirect assortative mating on genetic parameters following equations derived by Gianola (13). Specifically let *A* and *F* be two phenotypes in a population. Furthermore assume that under random mating in said population the heritabilities of *A* and *F* are 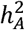 and 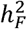 respectively and their genetic correlation is *ρ*_*g*_. Assortative mating on phenotype *A* leads to changes in the heritabilities of *A* and *F* as well as their genetic correlations, and under continued assortative mating these quantities will reach an equilibrium. Specifically under continued assortment on *A* with partner correlation *ρ*_*couple*_ the equilibrium heritability of the assortment trait *A* is given by

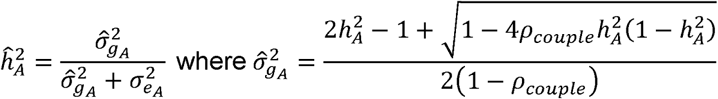

the equilibrium genetic correlation is given by

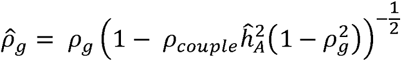

and the equilibrium heritability of the focal trait ***F*** given by

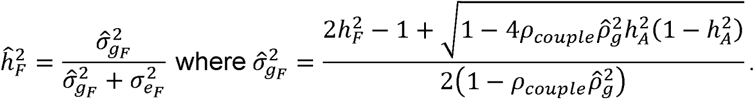

We provide an online calculator to compute the effects of direct and indirect assortative mating on genetic parameters (http://www.dissect.ed.ac.uk/projects/assortativemating.html).

### UK Biobank Couples

Identification of heterosexual couples in the UK Biobank has been previously reported (4). Specifically, using household sharing information we identified a set of 105,380 households with exactly two members in the cohort. Of these 90,297 satisfied all of the following criteria a) individuals reported different ages for one or both parents b) individuals had an age difference of less than 10 years c) individuals were of opposite gender d) both individuals reported to live only with their partner or partner and children. We restricted our analysis to a subset of 79,094 couples for which both partners self-reported to be of White-British ethnicity.

### UK Biobank Phenotypes

We utilized the family history for twelve diseases for both biological parents and age at death for both biological parents as provided by participants of the UK Biobank. Further information regarding these phenotypes can be obtained through the UK Biobank online documentation (http://biobank.ctsu.ox.ac.uk/crystal/index.cgi). To identify self-reported controls we utilized self-reported medical history following the methodology of Muñoz et al. (9) to match diseases to those reported for family history. We also used Birth Year and Townsend Deprivation Index as provided by the UK Biobank resource. The UK Biobank contains information about the coordinates of the birth location with a resolution of one kilometer (km). We excluded individuals with miscoded coordinates corresponding to birth in the Atlantic Ocean identified through visual inspection. As the resolution of the provided birth coordinates is too fine to allow for effective permutations, i.e., there are too few individuals sharing birth coordinates, we used a 15 km grid to define Birth Location. That is, we assign all individuals who share birth coordinates when divided by 15 km and rounded to an integer to the same Birth Location.

### FamiLinx Couples and Phenotypes

The FamiLinx cohort, consisting of 86,124,644 individuals, is based on publicly accessible genealogy data ranging back up to the early 15^th^ century and covering individuals born across the world, although individuals of European and North American birth dominate. Considering individuals with common offspring, we identified a set of 9,421,824 couples. In our analysis we restricted ourselves to a subset of individuals with full information regarding year of birth and death, latitude and longitude of the birth location. We removed individuals with a birth location along the zero meridian as visual inspection suggested majority of these to be coding errors. We furthermore removed individuals with lifespans below 30 or above 130 and those born before 1600, due to the sparsity and lower reliability of data before that date, and after 1910, due to the bias towards individuals with reduced lifespan after that date. Finally, we removed individuals who died during the American Civil War (year of death 1861 to 1865), the 1^st^ World War (year of death 1914 to 1918) and the 2^nd^ World War (year of death 1939 to 1945) due to the previously reported excess number of early death in these periods (15). This resulted in a dataset of 3,445,971 individuals containing 323,155 couples, 97,223 sets of fathers in law and 66,077 sets of mothers in law with lifespan information. To allow for effective permutation we defined a one degree latitude and longitude grid to define birth location. We computed adjusted lifespans as the difference between an individuals lifespan and the mean lifespan of the stratum defined by the individuals sex, birth year and birth location as defined above. As in the permutation analysis, we excluded all strata with fewer than 10 individuals.

### Estimation of genetic values

For our analysis, we used the data for the individuals genotyped in phase 1 of the UK Biobank genotyping program. 49,979 individuals were genotyped using the Affymetrix UK BiLEVE Axiom array and 102,750 individuals using the Affymetrix UK Biobank Axiom array. Details regarding genotyping procedure and genotype calling protocols are provided elsewhere (http://biobank.ctsu.ox.ac.uk/crystal/refer.cgi?id=155580). We performed quality control using the entire set of genotyped individuals before extracting the White-British cohort used in our analyses. From the overlapping markers between the two arrays, we excluded those which were multi-allelic, their overall missingness rate exceeded 2% or they exhibited a strong platform specific missingness bias (Fisher’s exact test, pval < 10^−100^). We also excluded individuals if they exhibited excess heterozygosity, as identified by UK Biobank internal QC procedures (http://biobank.ctsu.ox.ac.uk/crystal/refer.cgi?id=155580), if their missingness rate exceeded 5% or if their self-reported sex did not match genetic sex estimated from X chromosome inbreeding coefficients. These criteria resulted in a reduced dataset of 151,532 individuals. Finally, we only kept variants that did not exhibit departure from Hardy-Weinberg equilibrium (pval < 10^−50^) in the unrelated (i.e. with a relatedness below or equal to 0.0625) genotypically White-British subset of the cohort and had a MAF > 5%. To define the genotypically White-British subset, we performed a Principal Components Analysis (PCA) of all individuals passing genotypic QC using a linkage disequilibrium pruned set of 99,101 autosomal markers (http://biobank.ctsu.ox.ac.uk/crystal/refer.cgi?id=149744) that passed our SNP QC protocol. The genotypically White-British individuals were defined as those for whom the projections onto the leading twenty genomic principal components fell within three standard deviations of the mean and who self-reported their ethnicity as White-British. We furthermore pruned the set of genotypically White-British individuals removing one individual from pairs with relatedness above 0.0625 (corresponding to second degree cousins) to obtain a datasets of unrelated genotypically White-British individuals. Employing these individuals we jointly estimated heritabilities and SNP effects following the mixed model approach using the DISSECT tool (19). All models included the leading 20 genomic principal components as fixed effects. In addition, models used to estimate genetic values for self-reported disease also included Sex, Age and Townsend Deprivation Index as fixed effects. For family disease history traits we fitted models with only genomic principal components and models which also included Birth Year and Birth Location as categorical and Parent Age as continuous covariates. Using the estimated SNP effects we obtained genetic values (i.e. Best Linear Unbiased Predictors) for 10,160 White-British couples where both individuals had been genotyped and computed their Pearson’s correlation. We combined paternal and maternal estimates using the Olkin-Pratt fixed effect approach (22). For self-reported and family history of disease we transformed heritabilities to the liability scale using the sample specific prevalence.

### Polygenic Risk Score for Longevity

We computed polygenic risk scores based on variants recently reported in a GWAS of parental longevity in the UK Biobank (23). Specifically, the polygenic risk score for an individual was computed as sums of dosages weighted by reference allele effects for variants with reported associations. We computed separate risk scores for paternal and maternal longevity in both cases using all variants associated at with a pvalue<10^−6^ (see Joshi et al. Supplementary Table 1). We used the imputed genotypes released by the UK Biobank resulting in polygenic risk scores based on 16 and 10 variants for paternal and maternal longevity respectively. We then computed Pearson’s correlations between polygenic risk scores of couples.

### Permutation Analysis

We stratified couples based on the Birth Year and Birth Locations of both partners and permuted male partners within each strata. To allow for effective permutations we only included couples in strata of size larger than 10 in the analysis. For each permutation we computed *χ*^2^ statistics for family history and Pearson’s correlations for parental longevity. Empirical pvalues where then computed as the fraction of statistics exceeding the statistic computed without permutation, based on 10,000 permutations.

### Effect of Year of Birth proxy

The UK Biobank does not directly contain information regarding the year’s of birth of parents of participants. As such we used the participant’s year of birth as a proxy measure of the parent’s year of birth in permuation analyses for both longevity and disease. For a subset of parent’s, specifically parents who are still alive at recruitment of the participant, we can infer the parent’s year of birth from the date of recruitment and the parent’s age. The subset of parents who are still alive is relatively small, only 22% of fathers and 39% mothers respectively, and is complementary to the set of parents used in the analysis of longevity who were required to be deceased. While we can therefore not use the data in our main analysis, it allows us to evaluate the effect of using a proxy measure.

The correlation between offspring and parent year of birth is relatively high with ρ=0.78. For family history of disease we performed two additional permutation analyses. On the subset of parents with available year of birth, we permuted UK Biobank couples within the years of birth of their parents. That is, the offspring within the years of birth of the parents. We did not permute within both Birth Year and Birth Location strata due to the smaller sample size. The results of these permutation analyses, albeit with a much smaller sample size, are consistent with the results obtained with the proxy measure, suggesting that adjusting for Year of Birth of the children is an acceptable, albeit not perfect, proxy for Year of Birth of the parents (SI Appendix Table S9).

### Correlations in Family History of Disease

As disease history or status for an individual is a binary trait, Pearson’s correlations are not a suitable measure of correlation. Instead we computed polychoric correlations (24) using the R package polycor (25). In addition we assessed dependence between partner’s family histories using a *χ*^2^ test and by computing empirical mutual information (26). For mutual information we computed an empirical pvalue for departure from independence using permutations. That is, we computed empirical mutual information for 1000 datasets in which family history for the male partners had been permuted and compared them to the empirical mutual information on the observed data.

### Regression Analysis

We computed linear regression models, regressing parental longevity on Birth Year, Birth Location, as well as Townsend Deprivation Index and height, waist to hip ratio, BMI and smoking history in Pack Years, and the squares of these factors, of their children. Birth Year and Birth Location were coded as categorical variables while all other factors and their squares were included as continuous variables. Using the fitted models we computed residuals and correlations between couples using these residuals. Comparing these, we quantified the change in correlations due to inclusion of individual covariates in the models.

## Acknowledgements

This work was mainly supported by The Roslin Institute Strategic Grant funding from the BBSRC. AT also acknowledges funding from the Medical Research Council Human Genetics Unit. This work used the ARCHER UK National Supercomputing Service (http://www.archer.ac.uk) and the Edinburgh Compute and Data Facility (ECDF) (http://www.ecdf.ed.ac.uk/). This research has been conducted using the UK Biobank Resource under project 6684.

## SI Appendix

**Figure S1:**
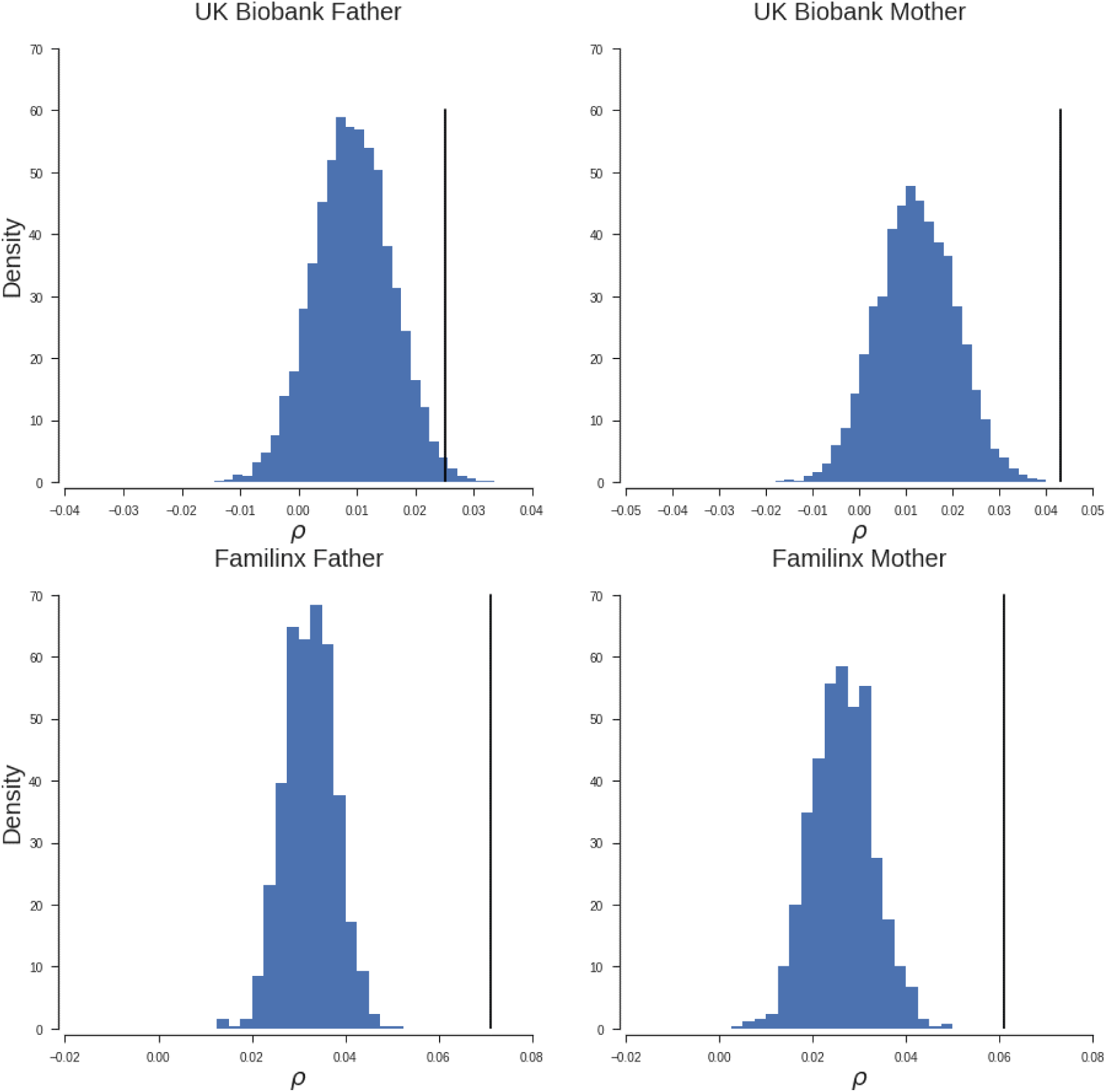
Density of correlations between parent’s life spans for 10,000 fictitious couples with assortment structure due to Birth Location and Birth Year matching that of observed couples in either the UK Biobank or Familinx cohort. The black vertical line indicates the correlations observed in real couples in the respective cohort.

**Table S1:** Phenotypic correlations between partners for potential explanatory variables.

|  |  | Male Partner |  |  |  |  |  |
| --- | --- | --- | --- | --- | --- | --- | --- |
|  |  | Birth Year | Townsend | Height | WHR | BMI | Pack Years |
| Female Partner | Birth Year | **0.91** | 0.04 | 0.14 | -0.14 | -0.01 | -0.17 |
|  | Townsend | 0.04 | **1.00** | -0.07 | 0.08 | 0.07 | 0.14 |
|  | Height | 0.15 | -0.06 | **0.26** | -0.06 | -0.04 | -0.07 |
|  | WHR | -0.15 | 0.09 | -0.06 | **0.19** | 0.16 | 0.11 |
|  | BMI | -0.07 | 0.09 | -0.07 | 0.22 | **0.24** | 0.08 |
|  | Pack Years | -0.10 | 0.17 | -0.06 | 0.12 | 0.09 | **0.34** |

Townsend = Townsend Deprivation Index, WHR = Waist to Hip ratio, BMI = Body Mass Index

**Table S2:** Residual partner correlations in different models of parental longevity.

| Model | N | Father |  |  | N | Mother |  |  |
| --- | --- | --- | --- | --- | --- | --- | --- | --- |
|  |  | r | p | % |  | r | p | % |
| Null | 22824 | 0.025 | 0.01 | 1.00 | 15026 | 0.043 | 2×10<sup>-4</sup> | 1.00 |
| Birth Info. | 22824 | 0.012 | 0.21 | 0.47 | 15026 | 0.019 | 0.10 | 0.44 |
| All | 22824 | 0.006 | 0.52 | 0.25 | 15026 | 0.013 | 0.24 | 0.32 |
| **Individual Factors** |  |  |  |  |  |  |  |  |
| BMI | 22824 | 0.010 | 0.28 | 0.40 | 15026 | 0.017 | 0.13 | 0.41 |
| WHR | 22824 | 0.010 | 0.28 | 0.41 | 15026 | 0.017 | 0.14 | 0.40 |
| Height | 22824 | 0.012 | 0.21 | 0.47 | 15026 | 0.018 | 0.11 | 0.43 |
| Townsend | 22824 | 0.009 | 0.32 | 0.38 | 15026 | 0.017 | 0.14 | 0.39 |
| Pack Years | 22824 | 0.008 | 0.37 | 0.34 | 15026 | 0.017 | 0.14 | 0.39 |

Null = no covariates, Birth Info.= Birth Year and Birth Location, All = Birth Info and all considered covariates and their squares, Models for individual covariates, contain Birth Info., the covariate and the covariate squared, N = number of couples, $\rho$ = Pearson's correlation of residuals, $P$ = pvalue for $\rho = 0$ , % = fraction of $\rho$ under null model remaining

**Table S3:** Estimates of heritability for parental longevity.

|  | N | h<sup>2</sup> |  |
| --- | --- | --- | --- |
|  |  | estimate | s.e. |
| Father | 79216 | 0.04 | 0.005 |
| Mother | 64002 | 0.03 | 0.006 |

N = number of individuals used in fitting model, h<sup>2</sup> = heritability

**Table S4:** Measures of association of family history in all couples.

|  |  | **N** | **MI** | **$P_{MI}$** | **$P_{\chi^2}$** | **$P_{\rho}$** |
| --- | --- | --- | --- | --- | --- | --- |
| **Father** | **Heart disease** | 69751 | 0.0003 | <0.001 | $1 \times 10^{-10}$ | $6 \times 10^{-11}$ |
|  | **Stroke** | 69751 | 0.00006 | <0.001 | 0.0051 | 0.003 |
| | **Lung cancer** | 68129 | 0.0001 | <0.001 | 0.0002 | $1 \times 10^{-4}$ |
|  | **Bowel cancer** | 68129 | 0.00004 | 0.03 | 0.0169 | 0.009 |
| | **Chronic bronchitis/emphysema** | 69751 | 0.00025 | <0.001 | $2 \times 10^{-9}$ | $2 \times 10^{-9}$ |
| | **High blood pressure** | 69751 | 0.0011 | <0.001 | $5 \times 10^{-36}$ | $1 \times 10^{-35}$ |
|  | **Diabetes** | 69751 | 0.00002 | 0.11 | 0.1366 | 0.067 |
| | **Alzheimer's disease/dementia** | 69751 | 0.00012 | <0.001 | $2 \times 10^{-5}$ | $2 \times 10^{-5}$ |
| | **Parkinson's disease** | 68129 | $3 \times 10^{-10}$ | 0.5 | 0.5823 | 0.267 |
|  | **Severe depression** | 68129 | 0.00001 | 0.13 | 0.2193 | 0.103 |
|  | **Prostate cancer** | 68129 | 0.00005 | <0.001 | 0.0073 | 0.004 |
| **Mother** | **Heart disease** | 73308 | 0.00065 | <0.001 | $7 \times 10^{-23}$ | $9 \times 10^{-23}$ |
| | **Stroke** | 73308 | 0.0003 | <0.001 | $2 \times 10^{-11}$ | $2 \times 10^{-11}$ |
| | **Lung cancer** | 72160 | 0.00012 | <0.001 | $1 \times 10^{-5}$ | $1 \times 10^{-5}$ |
| | **Bowel cancer** | 72160 | $3 \times 10^{-10}$ | 0.57 | 0.533 | 0.747 |
| | **Breast cancer** | 72160 | $1 \times 10^{-10}$ | 0.64 | 0.667 | 0.325 |
| | **Chronic bronchitis/emphysema** | 73308 | 0.0001 | <0.001 | $1 \times 10^{-4}$ | $7 \times 10^{-5}$ |
| | **High blood pressure** | 73308 | 0.00111 | <0.001 | $1 \times 10^{-37}$ | $7 \times 10^{-38}$ |
|  | **Diabetes** | 73308 | 0.00007 | <0.001 | 0.001 | 0.001 |
| | **Alzheimer's disease/dementia** | 73308 | 0.00035 | <0.001 | $1 \times 10^{-13}$ | $3 \times 10^{-13}$ |
|  | **Parkinson's disease** | 72160 | 0.00001 | 0.33 | 0.297 | 0.13 |
|  | **Severe depression** | 72160 | 0.00005 | <0.001 | 0.009 | 0.005 |

N=Number of Couples, MI = Mutual Information, $P_{MI}$ = empirical permutation based P value based on MI, $P_{\chi^2}$ = Chi Squared test P value, $P_{\rho}$ = P value for polychoric correlations

**Table S5:** Measures of association of family history in control couples.

|  |  | **N** | **MI** | **$P_{MI}$** | **$P_{\chi^2}$** |
| --- | --- | --- | --- | --- | --- |
| **Father** | **Heart disease** | 47786 | 0.00019 | <0.001 | $2 \times 10^{-5}$ |
|  | **Stroke** | 56239 | 0.00004 | 0.01 | 0.037 |
|  | **Lung cancer** | 57571 | 0.00009 | <0.001 | 0.002 |
|  | **Bowel cancer** | 56869 | 0.00006 | 0.01 | 0.009 |
| | **Chronic bronchitis/emphysema** | 55905 | 0.00020 | <0.001 | $2 \times 10^{-6}$ |
| | **High blood pressure** | 29573 | 0.00151 | <0.001 | $7 \times 10^{-22}$ |
| | **Diabetes** | 52964 | $2 \times 10^{-7}$ | 0.76 | 0.907 |
| | **Alzheimer's disease/dementia** | 58008 | 0.00012 | <0.001 | $1 \times 10^{-4}$ |
| | **Parkinson's disease** | 57408 | $1 \times 10^{-6}$ | 0.71 | 0.741 |
|  | **Severe depression** | 50056 | 0.00002 | 0.17 | 0.181 |
|  | **Prostate cancer** | 56596 | 0.00007 | <0.001 | 0.006 |
| **Mother** | **Heart disease** | 47786 | 0.00053 | <0.001 | $8 \times 10^{-13}$ |
| | **Stroke** | 56239 | 0.00029 | <0.001 | $6 \times 10^{-9}$ |
| | **Lung cancer** | 57571 | 0.00014 | <0.001 | $3 \times 10^{-5}$ |
| | **Bowel cancer** | 56869 | $4 \times 10^{-6}$ | 0.45 | 0.514 |
| | **Breast cancer** | 55075 | $3 \times 10^{-7}$ | 0.81 | 0.868 |
|  | **Chronic bronchitis/emphysema** | 55905 | 0.00007 | <0.001 | 0.003 |
| | **High blood pressure** | 29573 | 0.00096 | <0.001 | $4 \times 10^{-14}$ |
|  | **Diabetes** | 52964 | 0.00003 | 0.09 | 0.091 |
| | **Alzheimer's disease/dementia** | 58008 | 0.00038 | <0.001 | $7 \times 10^{-12}$ |
|  | **Parkinson's disease** | 57408 | 0.00001 | 0.32 | 0.331 |
|  | **Severe depression** | 50056 | 0.00005 | <0.001 | 0.025 |

N=Number of Couples, MI = Mutual Information, $P_{MI}$ = empirical permutation based P value based on MI, $P_{\chi^2}$ = Chi Squared test P value

**Table S6:** Empirical P values for association of family history based on permutations.

|  | **Father** | **Mother** |
| --- | --- | --- |
| **Alzheimer's disease/dementia** | 0.0026 | <0.0001 |
| **Bowel cancer** | 0.0408 | 0.8155 |
| **Breast cancer** | - | 0.2781 |
| **Chronic bronchitis/emphysema** | <0.0001 | 0.0041 |
| **Diabetes** | 0.8072 | 0.1212 |
| **Heart disease** | <0.0001 | <0.0001 |
| **High blood pressure** | <0.0001 | <0.0001 |
| **Lung cancer** | 0.0316 | 0.0002 |
| **Parkinson's disease** | 0.8469 | 0.3366 |
| **Prostate cancer** | 0.0149 | - |
| **Severe depression** | 0.0732 | 0.0344 |
| **Stroke** | 0.0441 | 0.0074 |

**Table S7:** Estimates of heritability for family history traits.

| | | Controls | Cases | $h^2$ | | $h_e^2$ |
| --- | --- | --- | --- | --- | --- | --- |
|  |  |  |  | est. | s.e. |  |
| Father | Heart disease | 69,745 | 31,053 | 0.033 | 0.004 | 0.05 |
|  | Stroke | 86,219 | 14,579 | 0.005 | 0.0037 | 0.01 |
|  | Lung cancer | 90,372 | 9,088 | 0.009 | 0.0034 | 0.03 |
|  | Bowel cancer | 93,880 | 5,580 | 0.012 | 0.0038 | 0.05 |
|  | Breast cancer | - | - | - | - | - |
|  | Chronic bronchitis/emphysema | 89,434 | 11,364 | 0.022 | 0.0039 | 0.06 |
|  | High blood pressure | 79,773 | 21,025 | 0.024 | 0.004 | 0.04 |
|  | Diabetes | 91,804 | 8,994 | 0.026 | 0.004 | 0.08 |
|  | Alzheimer's disease/dementia | 96,295 | 4,503 | 0.008 | 0.0036 | 0.04 |
|  | Parkinson's disease | 97,142 | 2,318 | 0.003 | 0.0035 | 0.02 |
|  | Severe depression | 95,986 | 3,474 | 0.008 | 0.0035 | 0.04 |
|  | Prostate cancer | 92,526 | 6,934 | 0.001 | 0.0028 | 0.005 |
| Mother | Heart disease | 83,680 | 20,117 | 0.016 | 0.0036 | 0.03 |
|  | Stroke | 89,469 | 14,328 | 0.006 | 0.0034 | 0.01 |
|  | Lung cancer | 98,437 | 4,366 | 0.008 | 0.0034 | 0.04 |
|  | Bowel cancer | 97,576 | 5,227 | 0.012 | 0.0037 | 0.05 |
|  | Breast cancer | 94,606 | 8,197 | 0.019 | 0.0038 | 0.06 |
|  | Chronic bronchitis/emphysema | 97,417 | 6,380 | 0.014 | 0.0037 | 0.05 |
|  | High blood pressure | 73,402 | 30,395 | 0.031 | 0.0039 | 0.05 |
|  | Diabetes | 94,316 | 9,481 | 0.03 | 0.0039 | 0.09 |
|  | Alzheimer's disease/dementia | 95,336 | 8,461 | 0.022 | 0.0038 | 0.07 |
|  | Parkinson's disease | 101,229 | 1,574 | 0.001 | 0.0033 | 0.01 |
|  | Severe depression | 96,333 | 6,470 | 0.009 | 0.0037 | 0.03 |
|  | Prostate cancer | - | - | - | - | - |

Controls/Cases= number of controls and cases used to fit the model, $h^2$ = heritability on the observed scale, $h_e^2$ = heritability on the liability scale

**Table S8:** Within couple correlations of genetic values for family history and self-reported disease with adjustment for Birth Year, Birth Location and Parent Age.

|  | Father |  | Mother |  | Combined<sup>†</sup> |  |
| --- | --- | --- | --- | --- | --- | --- |
| | $\rho$ | $P$ | $\rho$ | $P$ | $\rho$ | $P$ |
| Heart disease | 0.011 | 0.28 | 0.016 | 0.11 | 0.013 | 0.03 |
| Stroke | 0 | 0.97 | 0.02 | 0.05 | 0.010 | 0.17 |
| Lung cancer | -0.007 | 0.47 | 0.012 | 0.22 | 0.002 | 0.40 |
| Bowel cancer | -0.001 | 0.93 | -0.016 | 0.11 | -0.008 | 0.13 |
| Breast cancer | - | - | -0.003 | 0.77 | - | - |
| Chronic bronchitis/emphysema | -0.003 | 0.79 | 0.03 | 0.003 | 0.014 | 0.20 |
| High blood pressure | 0.023 | 0.02 | 0.028 | 0.005 | 0.025 | $1.7 \times 10^{-4}$ |
| Diabetes | 0.004 | 0.7 | 0.008 | 0.41 | 0.006 | 0.20 |
| Alzheimer's disease/dementia | -0.002 | 0.86 | 0.002 | 0.83 | 0.000 | 0.49 |
| Parkinson's disease | -0.001 | 0.89 | -0.001 | 0.93 | -0.001 | 0.43 |
| Severe depression | 0.015 | 0.12 | -0.008 | 0.43 | 0.004 | 0.37 |
| Prostate cancer | 0.007 | 0.46 | - | - | - | - |

<sup>†</sup>meta-analysis of paternal and maternal correlations, $\rho$ = Pearson's correlation of residuals, $P$ = pvalue for $\rho=0$

**Table S9:**
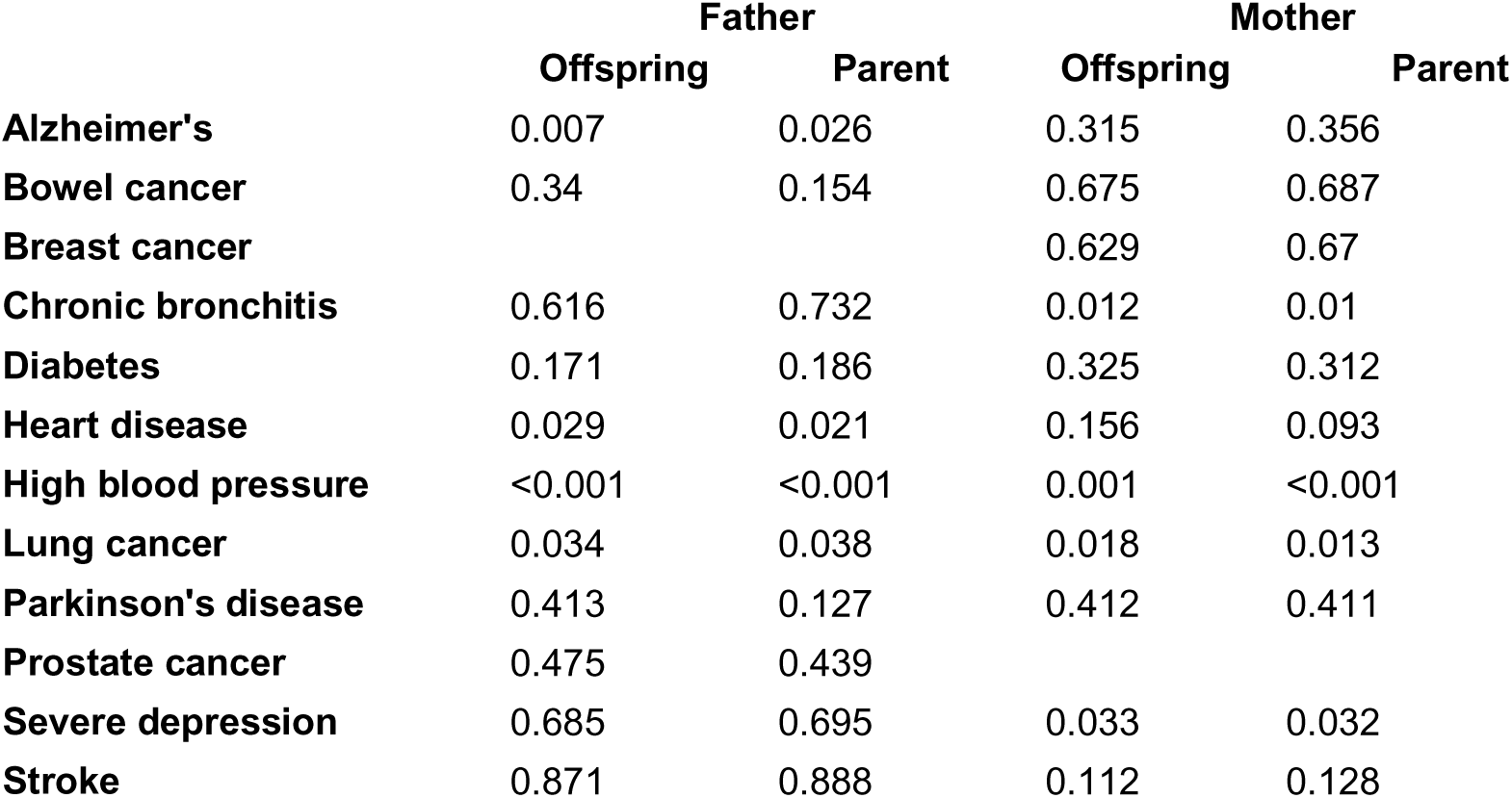
Empirical P values from permutation test within offspring’s and parent’s year of birth strata.

|  | Father |  | Mother |  |
| --- | --- | --- | --- | --- |
|  | Offspring | Parent | Offspring | Parent |
| Alzheimer's | 0.007 | 0.026 | 0.315 | 0.356 |
| Bowel cancer | 0.34 | 0.154 | 0.675 | 0.687 |
| Breast cancer |  |  | 0.629 | 0.67 |
| Chronic bronchitis | 0.616 | 0.732 | 0.012 | 0.01 |
| Diabetes | 0.171 | 0.186 | 0.325 | 0.312 |
| Heart disease | 0.029 | 0.021 | 0.156 | 0.093 |
| High blood pressure | <0.001 | <0.001 | 0.001 | <0.001 |
| Lung cancer | 0.034 | 0.038 | 0.018 | 0.013 |
| Parkinson's disease | 0.413 | 0.127 | 0.412 | 0.411 |
| Prostate cancer | 0.475 | 0.439 |  |  |
| Severe depression | 0.685 | 0.695 | 0.033 | 0.032 |
| Stroke | 0.871 | 0.888 | 0.112 | 0.128 |

